# Valency-Limited Molecular Dynamics Simulations of Stickers-and-Spacers Polymers Reveal a Tradeoff Between Condensation and Organization

**DOI:** 10.64898/2026.08.02.742325

**Authors:** Yumeng Zhang, Amogh Sood, Advait Athreya, Bin Zhang

## Abstract

Biomolecular condensates formed by intrinsically disordered proteins (IDPs) are often described using stickers-and-spacers models, in which specific sticker motifs form reversible crosslinks and spacer regions modulate phase behavior. Recent theory predicts that heterogeneous nonspecific spacer interactions can promote condensation but may also disrupt sticker-mediated organization. Here, we develop an off-lattice coarse-grained stickers-and-spacers polymer model for continuous molecular dynamics simulations and implement it in the GPU-accelerated OpenABC package. The model uses a directional sticker–sticker interaction to encode limited valency through interaction geometry, producing effectively one-to-one sticker binding without explicit bond assignment. Simulations of one-component systems show that sticker affinity and multivalency promote porous, network-like condensates, while nonspecific spacer interactions can also drive phase separation but produce more compact, spacer-dominated dense phases. When both interaction types are present, strong spacer heterogeneity reduces sticker conversion, suppresses sticker mobility, and disrupts the sticker-mediated network. In two-component systems, specific sticker interactions buffer client recruitment into host condensates, while nonspecific spacer interactions produce reservoir-dependent uptake. These results support a tradeoff in which spacer heterogeneity promotes condensation at the cost of condensate organization and compositional robustness, providing a physical rationale for the suppression of promiscuous spacer interactions in low-complexity IDP regions.

## Introduction

Biomolecular condensates organize cellular chemistry by selectively concentrating proteins, nucleic acids, and other biomolecules in membraneless compartments.^1–5^ By creating local environments distinct from the surrounding nucleoplasm or cytoplasm, they support processes including RNA splicing, ribosomal RNA processing, transcriptional regulation, and stress responses.^1,4,6,7^ Understanding how molecular sequence encodes condensate stability, internal organization, and molecular composition is therefore essential for explaining how condensates achieve biological specificity, but remains a central challenge in condensate biophysics.^4,8,9^

A major difficulty is the wide variety of chemical compositions and interaction motifs through which condensate-forming biomolecules, particularly intrinsically disordered proteins (IDPs), encode phase behavior.^8,10–12^ Experimental characterization and molecular simulations provide detailed insight into individual systems,^13–46^ but extracting general physical principles from these studies remains challenging. Simplified theoretical frameworks therefore play an important role.^4,9,47–60^ Among these, the stickers-and-spacers model has been especially useful.^61–66^ In this framework, proteins are represented as polymers containing specific interaction motifs, or stickers, connected by flexible spacer regions. This coarse-grained description connects condensate formation to the physics of associative polymers and provides a natural language for understanding coupled phase separation, percolation, and gelation.^62,67,68^

The stickers-and-spacers model also highlights a fundamental feature of many condensate-forming IDPs: strong, privileged interaction motifs are often embedded within low-complexity regions.^54,61,69,70^ This architecture differs from a homogeneous polymer model, such as a simple Flory–Huggins description, because sticker-mediated physical crosslinks can generate networked condensates with percolated or gel-like structures.^51,52,62,64,71^ Such behavior has been observed in both theoretical studies and experiments on associative biomacromolecules.^64,72^ However, the prevalence of sticker–spacer architectures raises an important question. If non-specific attractive interactions among many residues can promote phase separation, why do many IDPs contain low-complexity regions that restrict sequence diversity and potentially suppress promiscuous spacer-mediated interactions? This question is also practically important because identifying stickers from sequence alone remains difficult when both specific and nonspecific interactions contribute to condensation.^5,10,70^

Recently, Sood and Zhang ^66^ proposed the stickers and random spacers model, or STARS, to address this question. The model generalizes the stickers-and-spacers framework by incorporating heterogeneous, nonspecific interactions among spacer segments in addition to specific, valency-limited sticker interactions. In this formulation, sequence diversification is represented by increased heterogeneity in nonspecific spacer interaction strengths, whereas low-complexity spacers correspond to a reduced spectrum of such interactions.^63,68,73,74^ The theory predicts a tradeoff: heterogeneous spacer interactions can stabilize phase separation by providing additional nonspecific attractions, but they can also compete with sticker-mediated crosslinks and disrupt the well-defined network structure and composition of condensates. This prediction provides a possible physical rationale for the prevalence of low-complexity regions in IDPs. By suppressing spurious spacer interactions, low-complexity sequences may preserve the functional interaction networks formed by privileged sticker motifs.

The analytical treatment of the STARS model relies on mean-field approximations that simplify spatial correlations, chain connectivity, and the microscopic competition among interaction types. Direct numerical simulations are therefore needed to test whether the predicted tradeoff persists in an explicit polymer model. A key challenge is that theoretical stickers are typically assumed to form limited-valency physical crosslinks, whereas standard isotropic pair potentials do not naturally enforce this constraint in continuous molecular dynamics simulations.

Here, we develop an off-lattice coarse-grained stickers-and-spacers polymer model that implements the key features of the STARS framework in continuous molecular dynamics simulations. The model introduces a directional sticker–sticker interaction that encodes limited valency through interaction geometry rather than explicit bond assignment, and we implement it in the GPU-accelerated OpenABC package for efficient simulation of user-defined sticker–spacer sequences. Using this framework, we test how sticker interactions and nonspecific spacer interactions jointly regulate condensate formation, network organization, and molecular recruitment. Our simulations support the central STARS prediction that spacer heterogeneity can promote condensation while compromising sticker-mediated organization and compositional robustness, providing a physical rationale for the suppression of promiscuous spacer interactions in low-complexity IDP regions.

## Results

### A Valency-Limited Stickers-and-Spacers Model for Continuous Molecular Dynamics

To test the predictions of the STARS model in an explicit dynamical system, we developed an off-lattice, coarse-grained polymer model for stickers-and-spacers simulations. A central challenge in constructing such a model is enforcing the limited valency of sticker interactions. Existing simulation approaches often address this issue through Monte Carlo bonding or discrete partner-assignment schemes.^9,48,65,68,73,75^ Although such approaches can generate appropriate equilibrium ensembles, they do not provide continuous molecular dynamics trajectories of sticker binding and unbinding and are therefore less suited for studying condensate assembly kinetics.

We address this challenge by encoding sticker valency directly into the interaction geometry. Each polymer is represented as a coarse-grained backbone chain of spacer beads. At user-defined sticker positions, an auxiliary sticker bead is attached to the corresponding parent backbone bead, creating an oriented sticker site (Figure 1A). Sticker–sticker attraction is modeled using a directional, hydrogen-bond-like potential that depends on both the distance between the two sticker beads and the angular alignment of their parent–sticker vectors. The interaction is most favorable when the sticker beads are close and the two parent–sticker vectors are collinear (Figure 1C). Because this optimal geometry cannot be simultaneously satisfied by multiple binding partners, the potential strongly suppresses multivalent binding by a single sticker without requiring an explicit bond-assignment algorithm.

**Figure 1:**
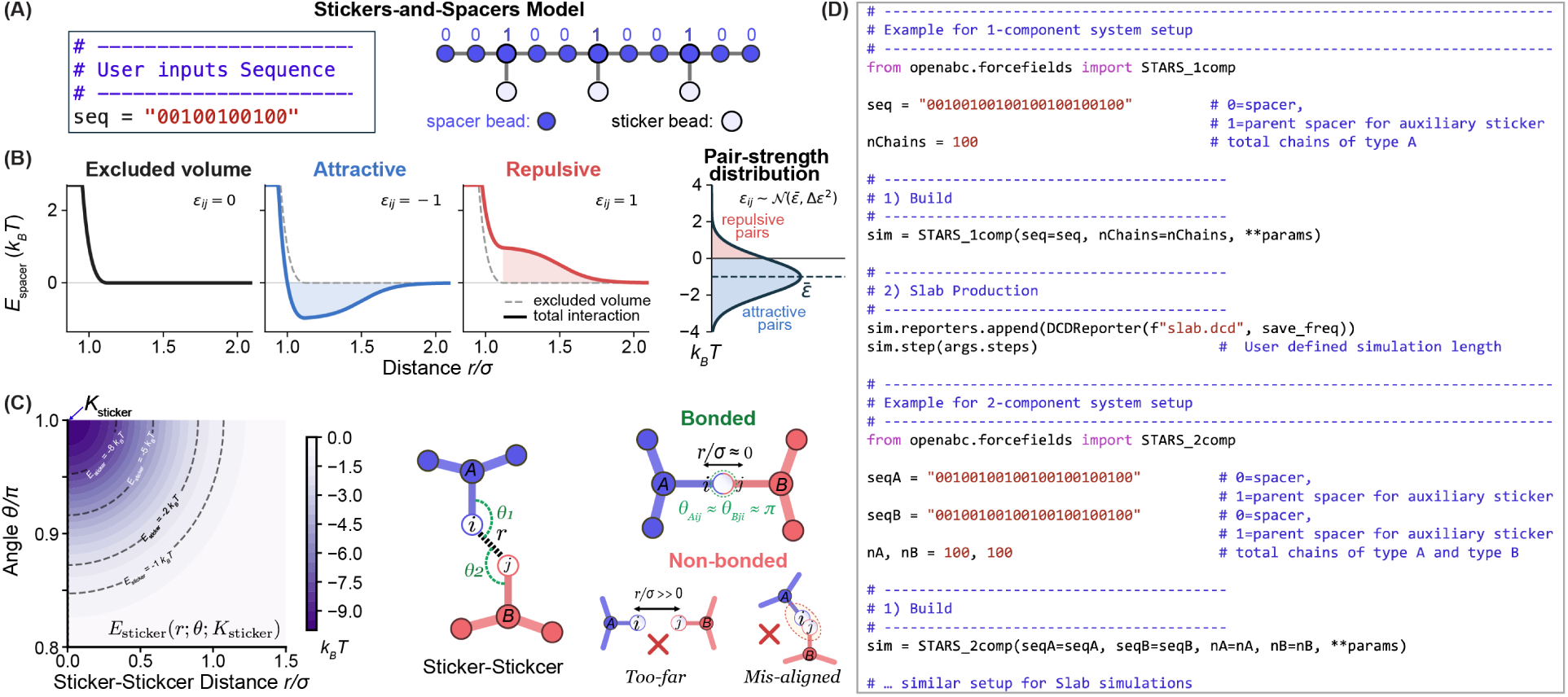
Definition and OpenABC implementation of a valency-limited stickers-and-spacers polymer model. (A) Polymer architectures are specified by binary sequences, where 0 denotes a pure spacer bead and 1 denotes a sticker-bearing spacer bead. Each sticker-bearing spacer is connected to an auxiliary sticker bead. (B) Illustration of spacer–spacer interaction potentials. In STARS simulations, the pairwise spacer interaction strength *ɛ_ij_* is sampled from a Gaussian distribution with mean *̅ɛ* and standard deviation Δ*ɛ*. (C) Illustration of the directional sticker–sticker interaction, which favors one-to-one sticker pairing and suppresses simultaneous binding to multiple partners. (D) Example Python scripts for setting up one-component and two-component polymer simulations in OpenABC.

The model also supports nonspecific spacer–spacer interactions (Figure 1B). Spacer beads interact through excluded-volume repulsion to prevent unphysical overlap, together with optional attractive or repulsive nonspecific interactions. For STARS simulations, pairwise spacer interaction strengths are sampled from a Gaussian distribution, allowing the mean spacer attraction and spacer-interaction heterogeneity to be controlled independently. The detailed functional forms of the interaction potentials are provided in the *Methods* section.

We implemented the polymer model in OpenABC,^76^ an OpenMM^77^-based, GPU-accelerated simulation platform for coarse-grained biomolecular condensate simulations. The implementation supports user-defined sticker–spacer architectures, homotypic and heterotypic polymer mixtures, independently tunable sticker–sticker and spacer–spacer interaction parameters, and arbitrary chain numbers. As illustrated in Figure 1D, OpenABC parses binary sequence specifications, constructs the corresponding polymer architectures, assigns the requested interaction parameters, and initializes condensate simulations through Python scripting. This implementation enables efficient long-timescale molecular dynamics simulations while leveraging existing OpenABC analysis tools to characterize phase separation, condensate organization, and condensate composition.

### Sticker-Mediated Condensation Produces Porous Network-like Condensates

Stickers-and-spacers polymers are associative polymers in which specific, reversible crosslinks can promote both condensation and network formation.^5,61–66,78,79^ In this framework, sticker-mediated contacts do more than provide cohesive energy: they also define the microscopic organization of the condensed phase. We therefore first asked whether our valency-limited off-lattice model captures the formation of porous, network-like condensates driven by reversible sticker–sticker interactions.

We simulated one-component systems containing 100 identical chains with 74 backbone spacer beads while varying the sticker–sticker interaction strength, *K*_sticker_. Each chain contained 24 auxiliary sticker beads, of which 12 were activated for sticker–sticker interactions; spacer–spacer attractions were turned off (Table S1). Weak sticker attractions did not support stable condensate formation (Figure 2A). As *K*_sticker_ became more attractive, polymers increasingly assembled into dense, slab-like structures, with robust condensation observed near *K*_sticker_ = −12 *k*_B_*T* and stronger assembly at larger sticker affinities. Even at *K*_sticker_ = −15 *k*_B_*T*, however, the condensed phase remained visibly porous, consistent with organization by an open sticker-contact network rather than nonspecific polymer collapse.

**Figure 2:**
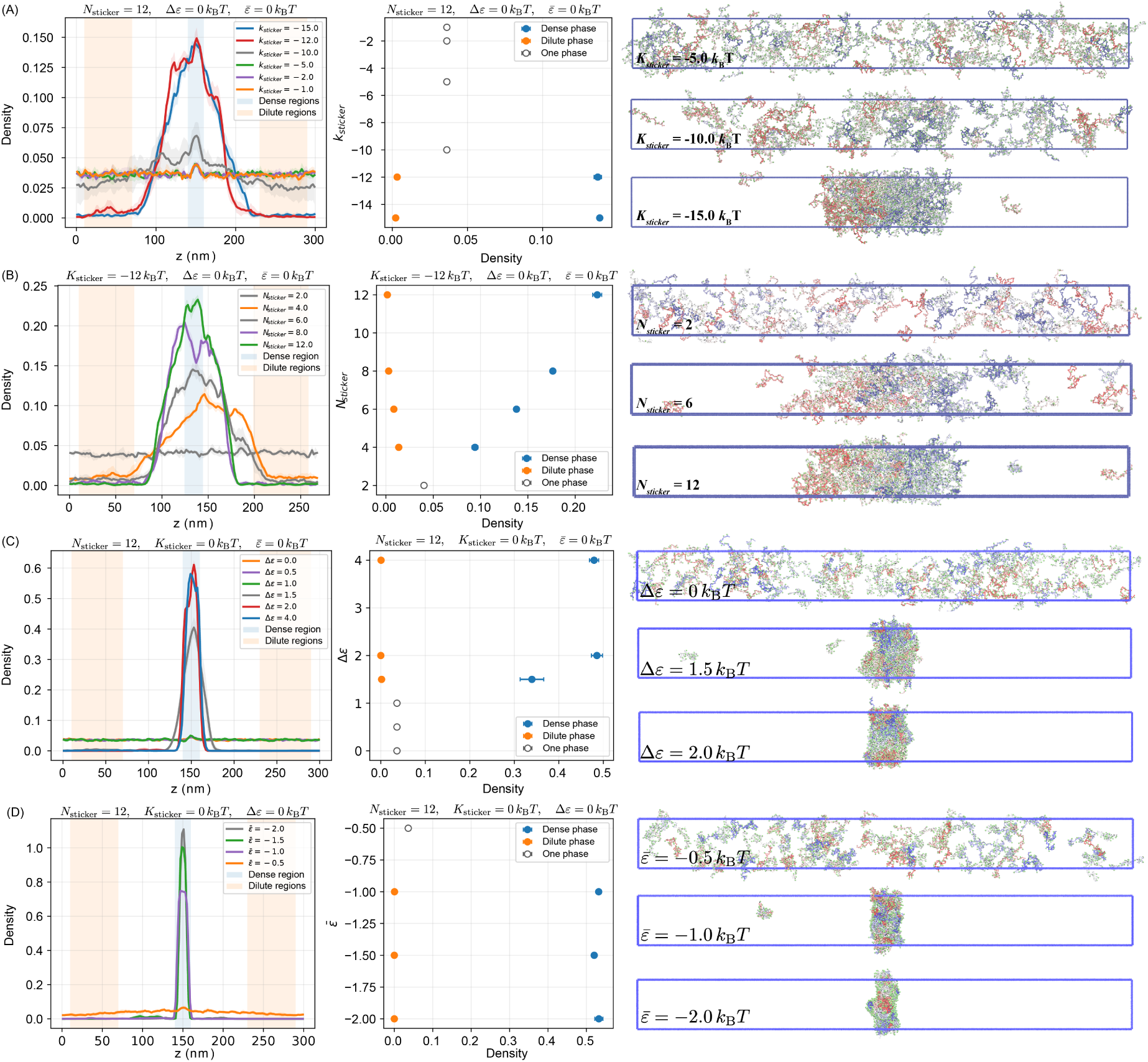
Sticker-mediated and spacer-mediated interactions produce condensates with distinct microscopic organization. (A) Increasing sticker affinity promotes the formation of porous, network-like condensates. Left: one-dimensional density profiles along the *z* axis. Blue shading marks the dense-phase region, and orange shading marks the dilute-phase windows used for density estimation. Middle: dense- and dilute-phase densities as a function of sticker interaction strength, *K*_sticker_. Right: representative simulation snapshots for selected *K*_sticker_ values. The blue outline denotes the simulation box, with the long axis corresponding to *z*. Coarse-grained polymer beads are colored by chain index, and sticker beads are highlighted as larger green spheres. (B) Effect of sticker multivalency, *N*_sticker_, at fixed sticker affinity. (C) Effect of spacer-interaction heterogeneity, Δ*ɛ*, in the absence of sticker attraction. (D) Effect of mean spacer interaction strength, *̅ɛ*, in the absence of sticker attraction.

We next examined sticker–sticker contacts directly to verify that the directional interaction enforces limited valency in dynamical trajectories. As expected, individual stickers formed at most one contact with another sticker under the conditions analyzed (Figures S1 and S3C), consistent with the intended valency-limited design. These contacts formed and dissociated reversibly during continuous molecular dynamics simulations, allowing sticker-network remodeling without imposing an explicit bond-assignment algorithm.

We then tested the role of sticker multivalency by varying the number of stickers per chain, *N*_sticker_, at fixed sticker affinity *K*_sticker_ = −12 *k*_B_*T* (Table S1). Chains with only two stickers did not form a stable condensate, whereas increasing *N*_sticker_ progressively enhanced condensation and produced well-defined dense phases at moderate multivalency (Figure 2B). These results are consistent with the associative-polymer picture in which both sticker affinity and sticker number regulate the formation of a connected sticker-contact network that stabilizes condensate assembly.

For comparison, we also simulated systems in which condensation was driven by nonspecific spacer–spacer interactions rather than sticker contacts. In these simulations, sticker–sticker attraction was eliminated by setting *K*_sticker_ = 0, while pairwise spacer interaction strengths were sampled from a Gaussian distribution with mean *̅ɛ* and standard deviation Δ*ɛ*. When Δ*ɛ* = 0, the model reduces to a homopolymer-like system with uniform spacer interactions. Making the mean spacer interaction *̅ɛ* more negative promoted condensation; however, the resulting dense phases were substantially more compact than the porous, sticker-mediated condensates observed in Figures 2A and 2B.

We further considered random-spacer systems with *̅ɛ* = 0. In this case, increasing Δ*ɛ* also promoted condensation (Figure 2C). A larger Δ*ɛ* broadens the distribution of spacer–spacer interaction strengths and increases the probability of strongly attractive spacer pairs, thereby stabilizing nonspecific polymer association even when the mean interaction strength is zero. Thus, both sticker-mediated crosslinking and nonspecific spacer heterogeneity can drive condensate formation, but they produce dense phases with distinct microscopic organization: sticker interactions favor porous, network-like condensates, whereas nonspecific spacer interactions favor more compact spacer-dominated assemblies.

### Nonspecific Spacer Interactions Stabilize Condensates but Disrupt Sticker Contacts

Having established that spacer heterogeneity can independently promote condensation, we next tested whether it competes with sticker-mediated organization, as predicted by the STARS theory.

We first established a sticker-only reference using the degree of sticker conversion, defined as the fraction of stickers participating in sticker–sticker contacts (see *Methods*). Because the directional potential limits each sticker to a single binding partner, the degree of conversion ranges from zero, when all stickers are unbound, to one, when all stickers participate in a contact. We further decomposed conversion into intrachain contacts, which form loops within individual polymers, and interchain contacts, which connect distinct polymers and contribute to network formation.

In the absence of spacer attractions, sticker conversion increased with stronger sticker affinity (Figure 3A). Increasing sticker affinity also shifted the contact distribution from intrachain loops toward interchain crosslinks, consistent with the formation of a more connected sticker-mediated network. This sticker-only reference established the expected dependence of network organization on specific sticker affinity.

**Figure 3:**
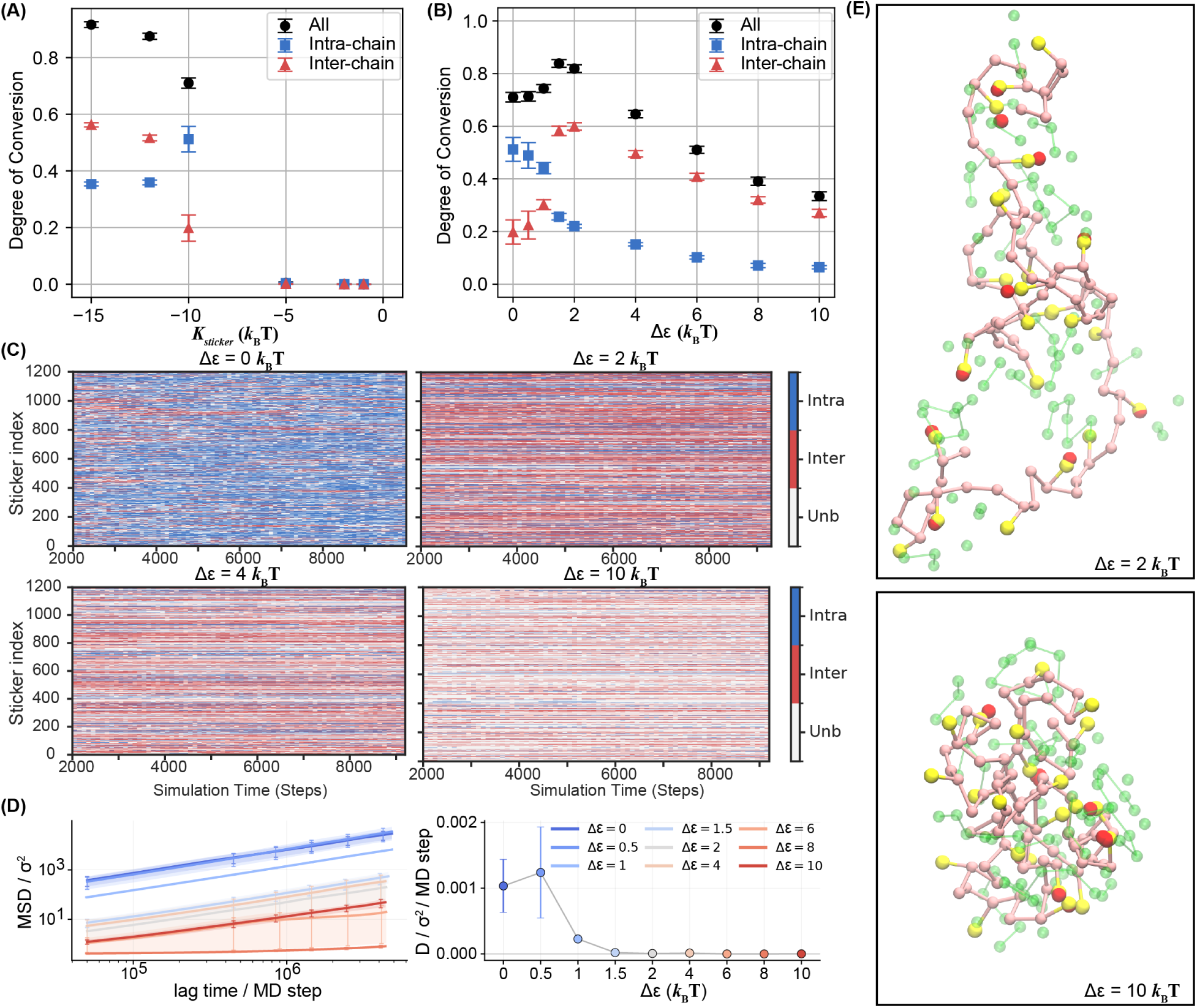
Spacer-interaction heterogeneity competes with sticker-mediated network organization. (A) Degree of sticker conversion as a function of sticker affinity *K*_sticker_ in the absence of spacer interactions. (B) Degree of sticker conversion as a function of spacer-interaction heterogeneity Δ*ɛ* at fixed sticker affinity *K*_sticker_ = −10 *k*_B_*T* . (C) Time-resolved sticker-contact states for representative Δ*ɛ* values. (D) Mean-squared displacement as a function of lag time and fitted diffusion coefficients of stickers at different Δ*ɛ* values. (E) Representative local snapshots for simulations with spacer-interaction heterogeneity Δ*ɛ* = 2 *k*_B_*T* and Δ*ɛ* = 10 *k*_B_*T* . A focal polymer chain is shown in pink with its stickers in yellow; neighboring spacers within 1.5*σ* and stickers within 0.5*σ* are shown in green and red, respectively.

We then fixed the sticker affinity at *K*_sticker_ = −10 *k*_B_*T*, near the onset of sticker-mediated phase separation. At this affinity, sticker contacts were present but not saturated, providing a sensitive regime for detecting either enhancement or disruption of sticker association by nonspecific spacer interactions. On top of the sticker interactions, we introduced nonspecific spacer–spacer interactions with strengths sampled from a Gaussian distribution with mean *̅ɛ* = 0. We then varied the standard deviation Δ*ɛ* to assess how spacer-interaction heterogeneity affects sticker-network organization.

At weak spacer heterogeneity, Δ*ɛ <* 2 *k*_B_*T*, the total degree of sticker conversion increased relative to the sticker-only case (Figure 3B). This increase was accompanied by a redistribution from intrachain to interchain contacts. This behavior is consistent with moderate spacer-mediated attractions increasing local polymer concentration and bringing stickers on different chains into closer proximity, thereby favoring intermolecular sticker contacts over intrachain loops.

At larger spacer heterogeneity, Δ*ɛ >* 2 *k*_B_*T*, both total and interchain sticker conversion decreased substantially (Figure 3B, Figure S2 and S3). Consistent with these ensemble averages, time-resolved contact-state analysis showed a larger fraction of unbound stickers at Δ*ɛ* = 4 and Δ*ɛ* = 10 throughout the simulations (Figure 3C). Thus, although stronger spacer heterogeneity provides an additional driving force for condensation, it reduces the fraction of stickers engaged in productive sticker–sticker contacts.

Representative local configurations illustrate this competition (Figure 3E). At Δ*ɛ* = 2, sticker contacts remain prominent around the focal polymer. At Δ*ɛ* = 10, the local environment is more compact and enriched in neighboring spacer segments, while fewer stickers participate in productive contacts. These snapshots suggest that strong spacer heterogeneity shifts local organization away from directional sticker-mediated binding and toward nonspecific spacer-dominated association.

Spacer-mediated compaction also reduced sticker mobility. Mean-squared displacement (MSD) analysis showed that increasing Δ*ɛ* decreased sticker diffusion within the condensate (Figure 3D). Reduced sticker mobility is expected to limit the ability of stickers to search for compatible binding partners and rearrange into the orientation required by the valency-limited interaction. Together, these results support the STARS prediction that spacer-interaction heterogeneity has opposing effects: moderate heterogeneity can stabilize condensates and promote interchain sticker engagement, whereas excessive heterogeneity disrupts sticker-mediated network organization by favoring nonspecific spacer-rich contacts.

### Limited-Valency Sticker Interactions Buffer Client Recruitment

Another key prediction of the STARS model is that recruitment mediated by specific, limited-valency sticker interactions should be more compositionally robust than recruitment driven by nonspecific spacer interactions. In particular, when client molecules are recruited through saturable sticker contacts, changes in client abundance inside the condensate are expected to be smaller than corresponding changes in the global client abundance. By contrast, nonspecific spacer-mediated interactions lack an intrinsic valency constraint and can therefore produce condensate compositions that are more sensitive to the external client reservoir.^80–86^

To test this prediction, we performed two sets of two-component simulations. In both cases, A molecules served as the host/scaffold species and formed condensates through A–A sticker interactions with *K*_AA_ = −15 *k*_B_*T* . We then compared two mechanisms of B-client recruitment. In the sticker-mediated recruitment model, A–B interactions were mediated by specific sticker–sticker attraction with *K*_AB_ = −17 *k*_B_*T*, and nonspecific spacer interactions were set to zero. In the spacer-mediated recruitment model, A–B sticker interactions were turned off by setting *K*_AB_ = 0, and A–B association was instead driven by nonspecific spacer interactions with *̅ɛ*_AB_ = 0 and Δ*ɛ*_AB_ = 1.5 *k*_B_*T* .

For both recruitment mechanisms, the number of host A molecules was held fixed while the total number of B molecules was varied from 100 to 300. Simulations were initialized from configurations in which A was packed into a condensate and B molecules were placed outside the condensate on one side of the slab. We then performed molecular dynamics simulations for 10^8^ steps and quantified B recruitment from the resulting density profiles. Additional details of the system setup and simulation protocol are provided in the *Methods* section.

The two recruitment mechanisms produced markedly different responses to increasing B abundance (Figure 4). When A–B interactions were mediated by specific stickers, increasing the total number of B molecules from 100 to 300 increased the number of recruited B molecules inside the A condensate only from 37 to 83 (Figure 4A, B). Thus, a threefold increase in the global B reservoir produced only a 2.2-fold increase in condensate-localized B. This sublinear response is consistent with saturable recruitment through a finite number of available A–B sticker contacts. Extending the analysis to *N*_B_ = 600 and varying the sticker–sticker interaction strength *K*_AB_ yielded consistent results (Figures S4 and S5).

**Figure 4:**
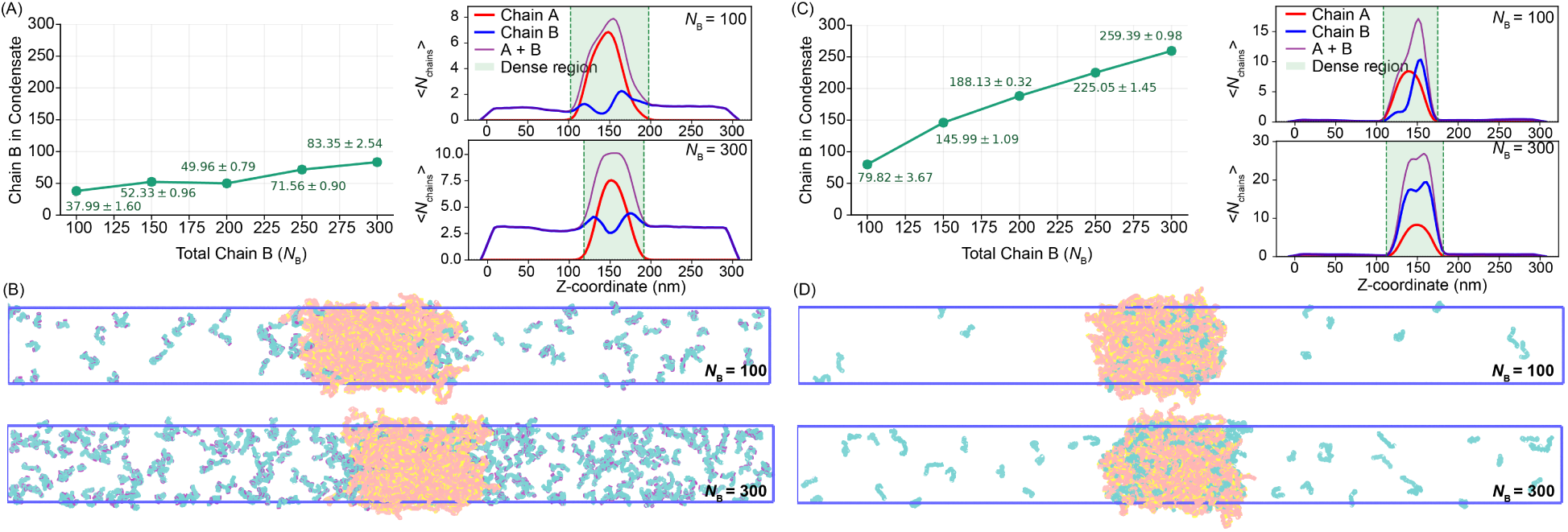
Specific sticker interactions buffer client recruitment in two-component condensates. (A) Number of B molecules recruited into the A condensate as a function of the total number of B molecules in the system for sticker-mediated A–B interactions. Density profiles are shown for representative simulations with *N*_B_ = 100 and *N*_B_ = 300. (B) Representative snapshots for the sticker-mediated condition at low (*N*_B_ = 100) and high (*N*_B_ = 300) B abundance. A and B chains are shown in red and green, respectively. A–A and A–B sticker types are shown in yellow and purple, respectively. (C) Number of B molecules recruited into the A condensate as a function of total B abundance for spacer-mediated A–B interactions. Density profiles are shown for representative simulations with *N*_B_ = 100 and *N*_B_ = 300. (D) Representative snapshots for the spacer-mediated condition at low (*N*_B_ = 100) and high (*N*_B_ = 300) B abundance.

In contrast, when A–B recruitment was mediated by nonspecific spacer interactions, B uptake was much more sensitive to the total number of B molecules. Under this condition, increasing the total number of B molecules from 100 to 300 increased the number of recruited B molecules inside the A condensate from 80 to 259 (Figure 4C, D, and Figure S6). Thus, spacer-mediated recruitment closely tracked the global B abundance, indicating that nonspecific interactions provide little buffering against changes in the client reservoir.

Together, these simulations support the STARS prediction that specific sticker interactions can impose compositional constraints on multicomponent condensates. Limited-valency A–B contacts produce saturable client recruitment and buffer condensate composition against changes in global client abundance. Nonspecific spacer interactions, by contrast, promote reservoir-dependent uptake and allow a wider range of condensate compositions. These results reinforce the central conclusion that sticker and spacer interactions can both promote co-condensation, but differ fundamentally in their ability to regulate condensate composition.

## Conclusions and Discussion

We developed and implemented in OpenABC an off-lattice molecular dynamics model that realizes effectively limited-valency sticker interactions through directional geometry. Using this implementation, we tested central predictions of the STARS theory. Our simulations support a tradeoff between condensation and organization: nonspecific spacer interactions can stabilize dense phases, but strong spacer heterogeneity disrupts sticker-mediated contacts, reduces sticker mobility, and promotes spacer-dominated condensate organization. In two-component systems, this distinction also affects molecular recruitment, with limited-valency sticker interactions buffering client uptake more effectively than nonspecific spacer interactions.

Although our simulations use simplified model systems, they suggest a physical rationale for the prevalence of low-complexity regions in phase-separating IDPs. If the only requirement were to promote phase separation, increasing spacer heterogeneity could be advantageous. However, functional condensates also require organized interaction networks and controlled molecular composition. Suppressing promiscuous spacer interactions through low-complexity sequence design may therefore help preserve functional sticker-mediated contacts, while conserved sticker motifs encode the specific interactions that define condensate organization and recruitment selectivity.

## Methods

### Energy Function of the Stickers-and-Spacers Polymer Model

We implemented a generalized stickers-and-spacers polymer model in OpenABC.^76^ The model is an off-lattice coarse-grained representation in which each polymer contains a backbone chain of spacer beads. A subset of backbone beads is designated as sticker-bearing, and each sticker-bearing backbone bead is connected to an auxiliary sticker bead (Figure 1A). Polymer architectures are specified by binary sequences, where 0 denotes a pure spacer bead and 1 denotes a sticker-bearing spacer bead. For example, the sequence 010 represents a three-bead backbone in which the central spacer bead carries an auxiliary sticker bead.

### The total potential energy is

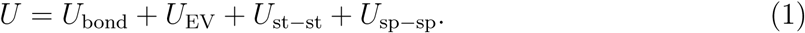

Here, *U*_bond_ maintains polymer connectivity, *U*_EV_ describes excluded-volume repulsion, *U*_st−st_ describes specific valency-limited sticker–sticker interactions, and *U*_sp−sp_ describes nonspecific spacer–spacer interactions. All energy and length parameters below are reported in reduced units of *ɛ* and *σ*, respectively.

### Bonded interactions

Bonded interactions were applied between adjacent backbone beads and between each auxiliary sticker bead and its parent backbone bead. These bonded pairs were connected using a class-2 bond potential,^87^

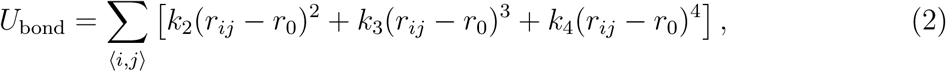

where *r_ij_* is the distance between bonded beads *i* and *j*, *r*_0_ is the equilibrium bond length, and the sum is over all bonded pairs. We used *r*_0_ = 1.0*σ*, *k*_2_ = 100 *ɛ/σ*^2^, *k*_3_ = 100 *ɛ/σ*^3^, and *k*_4_ = 100 *ɛ/σ*^4^.

### Excluded-volume interactions

Excluded-volume interactions were included to prevent unphysical overlap between nonbonded beads. We applied a purely repulsive Weeks–Chandler–Andersen (WCA) potential^88^ to nonbonded spacer–spacer and spacer–sticker pairs:

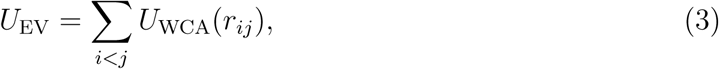

with

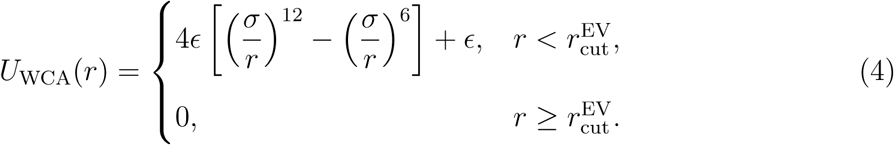

The cutoff was set to 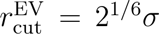, corresponding to the minimum of the Lennard-Jones potential. Directly bonded pairs, including adjacent backbone beads and each auxiliary sticker bead with its parent backbone bead, were excluded from *U*_EV_. Sticker–sticker pairs were also excluded from *U*_EV_, because their interactions were described separately by the orientation-dependent potential below.

### Sticker–sticker interactions

Specific sticker–sticker interactions were modeled using an orientation-dependent potential,

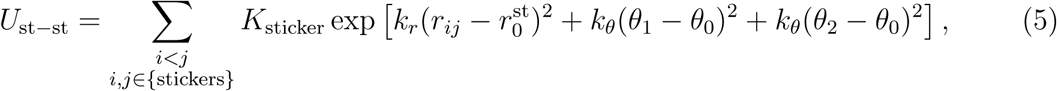

where *i* and *j* denote sticker beads, *r_ij_* is the sticker–sticker distance, and *θ*_1_ and *θ*_2_ are the two angles formed by the sticker beads and their corresponding parent backbone beads (Figure 1C). The interaction strength is controlled by *K*_sticker_, with more negative values corresponding to stronger attraction. We used *k_r_* = −2*σ*^−2^, *k_θ_* = −5, 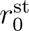, and *θ*_0_ = *π*.

With these parameters, the most favorable sticker–sticker interaction occurs when the two sticker beads overlap and the two parent–sticker vectors are collinear. This directional requirement encodes limited valency because a single sticker cannot simultaneously satisfy the optimal binding geometry with multiple partners. The functional form is analogous to orientation-dependent hydrogen-bond potentials used in coarse-grained molecular models.^89,90^

For computational efficiency, the sticker–sticker interaction was evaluated only within a finite radial cutoff,

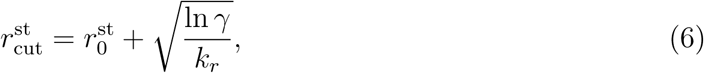

where *γ* = 10^−6^ specifies the value of the radial exponential factor at the cutoff.

### Spacer–spacer interactions

Nonspecific spacer–spacer interactions were modeled using a finite-range potential,^66^

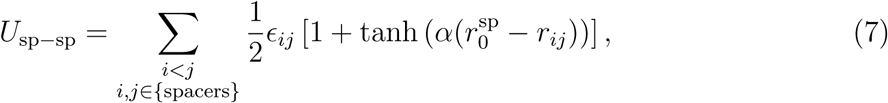

where *r_ij_* is the distance between spacer beads *i* and *j*, *α* controls the sharpness of the interaction, and 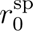 sets the midpoint of the interaction range. Negative values of *ɛ_ij_* correspond to attractive spacer–spacer interactions. We used *α* = 4.5*σ*^−1^ and 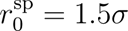 .

### The spacer–spacer interaction was truncated at

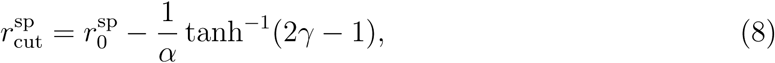

such that the prefactor 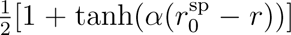 decays to *γ* = 10^−4^ at the cutoff. Bonded pairs were excluded from the spacer–spacer interaction.

### Molecular Dynamics Simulation Details

Molecular dynamics simulations were performed in OpenABC^76^ using OpenMM.^77^ The model was formulated in reduced units of mass *m*, length *σ*, and energy *ɛ*. To provide dimensional quantities required by OpenMM, we set *m* = 1 Da, *σ* = 1 nm, and *ɛ* = 1 kJ mol^−1^. Simulations were performed at *T* = *ɛ/k*_B_ ≈ 120.3 K, such that *k*_B_*T* = *ɛ*. This dimen-sionalization was introduced solely to express the reduced-unit model in units accepted by OpenMM; the resulting temperature and timescale are not intended as a direct mapping to a specific experimental system.

Initial configurations were generated by placing polymer chains in extended conformations on a three-dimensional grid. Chains were placed sequentially along the *x*, *y*, and *z* directions with a minimum separation of 5*σ* between neighboring chains. The initial placement was chosen to avoid steric overlap and to ensure that all chains were contained within the simulation box. For two-component systems, A chains were placed first, followed by B chains initialized at least 10*σ* away from the A-rich region to reduce initial A–B contacts.

After initial placement, each system was energy minimized and compressed under constant-pressure, constant-temperature (NPT) conditions. During this compression stage, only bonded and excluded-volume interactions were active for one-component systems, while sticker-sticker interactions were also included and set as −10*k_B_T* for two-component systems. Starting from the compressed configuration, the system was minimized again and relaxed in the constant-volume, constant-temperature (NVT) ensemble for 2 × 10^5^ steps using only bonded and excluded-volume interactions. The simulation box for slab simulations was then set to 30*σ* × 30*σ* × 300*σ*.

After this relaxation stage, each system was equilibrated for 5 × 10^3^ NVT steps using the full production potential energy. Production NVT simulations were then run for 5 × 10^7^ and 1 × 10^8^ steps for one-component and two-component simulations, respectively, with configurations saved every 5 × 10^3^ steps. Production simulations used Langevin dynamics with a friction coefficient of 0.1 ps^−1^ and a timestep of 1 fs.

### One-Component Systems

One-component simulations contained 100 identical A chains (Table S1). Each A chain contained 74 backbone spacer beads, of which 24 were sticker-bearing. Each sticker-bearing spacer was connected to one auxiliary sticker bead. For the one-component simulations described in the main text, only a subset of these sticker beads was activated for A–A sticker interactions, as specified in Table S1.

For each nonbonded spacer pair (*i, j*), the interaction strengths *ɛ_ij_* were sampled from Gaussian distributions, i.e.,

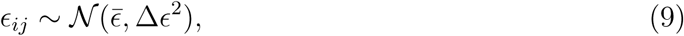

where *̅ɛ* controls the mean nonspecific spacer interaction and Δ*ɛ* controls the standard deviation of spacer-interaction heterogeneity.

### Two-Component Systems

Two-component simulations contained A and B chains with independently specified sticker–spacer architectures (Table S1). For A chains, 12 sticker beads were assigned to mediate A–A interactions and 12 sticker beads were assigned to mediate A–B interactions. B chains contained 14 backbone spacer beads and 4 auxiliary sticker beads assigned to A–B interactions. A–A and A–B sticker interactions were controlled independently through their corresponding sticker-interaction strengths, *K*_AA_ and *K*_AB_. Spacer–spacer interactions were similarly drawn from a normal distribution as in the one-component system.

### Data Analysis

#### Analysis of sticker valency

For a sticker *i*, we define its valency in time frame *t* as

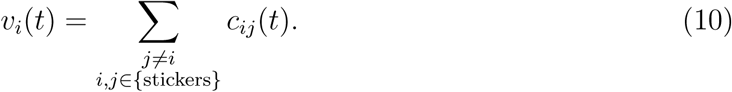

*c_ij_*(*t*) is a contact indicator function defined as

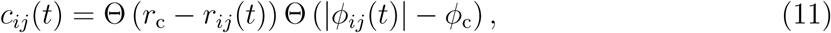

where *r*_c_ = 0.5*σ*, *φ*_c_ = 0.9*π*. *φ_ij_* is the dihedral angle defined by the four beads (*a, i, j, b*) in the range of (−*π, π*), where *a* and *b* correspond to the parent spacers connected to sticker *i* and *j*, respectively (Figure 1C).

The fractions of unbound, singly bound, and higher-valency stickers were calculated as the fractions of observations with *v_i_*(*t*) = 0, *v_i_*(*t*) = 1, and *v_i_*(*t*) *>* 1, respectively.

### Degree of conversion analysis

The degree of conversion was defined as

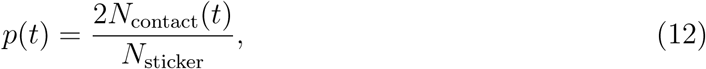

where *N*_sticker_ is the total number of sticker beads. The number of sticker–sticker contacts was calculated as

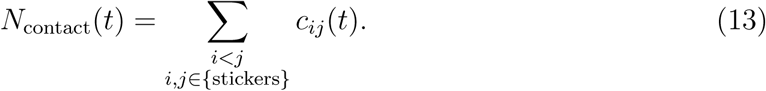

Here, *c_ij_*(*t*) is the contact indicator function defined in Eq. 11. Because higher-valency contacts were not observed under the analyzed conditions, this summation does not introduce double counting.

Contacts were further partitioned into intra-chain or inter-chain depending on whether the two stickers belonged to the same polymer chain. The corresponding conversion fractions were calculated as

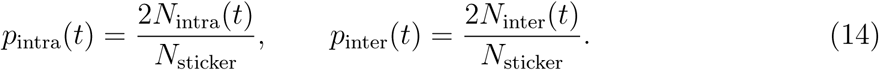

### Identify dense region in one/two-component systems

Dense-phase analysis was based on axial density profiles in both one- and two-component systems. The final 5 × 10^3^ saved frames of each production trajectory were used for analysis, from a total of 1 × 10^4^ and 2 × 10^4^ saved frames for the one- and two-component simulations, respectively. For each trajectory, these frames were divided into five contiguous blocks of equal length. Results were averaged within each block and then across the five blocks to obtain the mean, with uncertainties reported as the standard error of the mean across the five blockwise averages.

For one-component simulations, the aligned slab was centered in the simulation box, and the dense phase was evaluated within a fixed central core window of width 20 *σ* (blue regions in Figure 2). Two 60 *σ* dilute-phase windows, which are selected 10 *σ* away from the *z*-axis boundaries, were used as reference dilute regions (orange regions in Figure 2). A system was classified as phase separated when the core density satisfied *ρ*_core_*/ρ*_dilute_ ≥ 2.

For two-component systems, A molecules remained condensed under all conditions analyzed. We therefore used the A-chain density profile, *ρ*_A_(*z*), to define the spatial boundary of the condensate. For each frame, the dense-phase interval was identified as the contiguous region along *z* where *ρ*_A_(*z*) ≥ *ρ*_cut_, with *ρ*_cut_ set to 1% of the maximum A density in that frame. A and B chains were assigned to the dense phase if the mean *z* coordinate of their beads fell within this interval.

### Mean-squared-displacement analysis

Sticker mobility was quantified from the mean-squared displacement (MSD). For a lag time *τ*, the MSD was calculated by averaging over all selected sticker beads and all valid time origins,

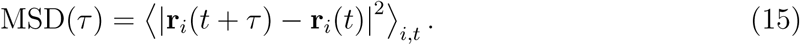

For each trajectory, the coordinates were unwrapped across periodic boundaries, and the first 20% of the trajectory was discarded as equilibration. Collective translational motion was removed by subtracting the displacement of the mean position of the selected sticker population. This correction removes drift of the sticker population as a whole, thereby isolating the relative mobility of individual stickers within the condensate.

The remaining trajectory was divided into five contiguous blocks of equal length. Within each block, MSD values were evaluated at up to 70 logarithmically spaced lag times extending to one half of the block duration. The MSD curves in Figure 3D show the mean across blocks, with uncertainties given as the standard error of the mean across blockwise MSD values.

The long-time diffusion coefficient was obtained by independently fitting the MSD in each block to

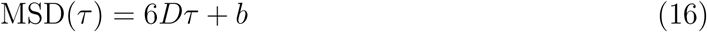

over lag times between 30% and 90% of the maximum sampled lag time. The diffusion coefficient reported in Figure 3D is the mean of the five blockwise fitted values, and its uncertainty is the standard error of the mean across blocks. Lag times are reported in MD steps, MSD values are reported in units of *σ*^2^, and diffusion coefficients are reported in units of *σ*^2^ per MD step.

## Acknowledgement

This work was supported by the National Institutes of Health (Grant R35GM133580).

## Data and materials availability

The stickers and spacers polymer model is implemented in the open source software package OpenABC. Tutorials for setting up simulations can be found on GitHub: STARS.

