## Supplemental Material for "Valency-Limited Molecular Dynamics Simulations of Stickers-and-Spacers Polymers Reveal a Tradeoff Between Condensation and Organization"

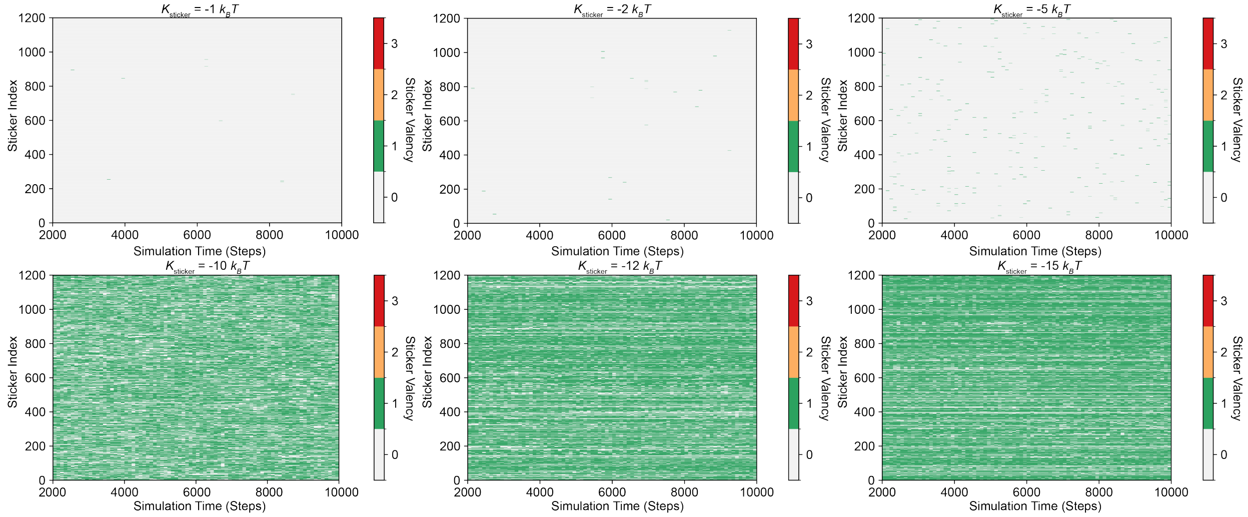


Figure S1: **Time-resolved sticker valency at different sticker interaction strengths.** Each row corresponds to an individual sticker, indexed on the $y$-axis, and the $x$-axis denotes simulation time. Color indicates the instantaneous sticker valency, $v_i(t)$, defined as the number of other stickers bound to sticker $i$ at time $t$ as in Eq. 10 of the main text. The trajectories are the same as those analyzed in Figure 3A.

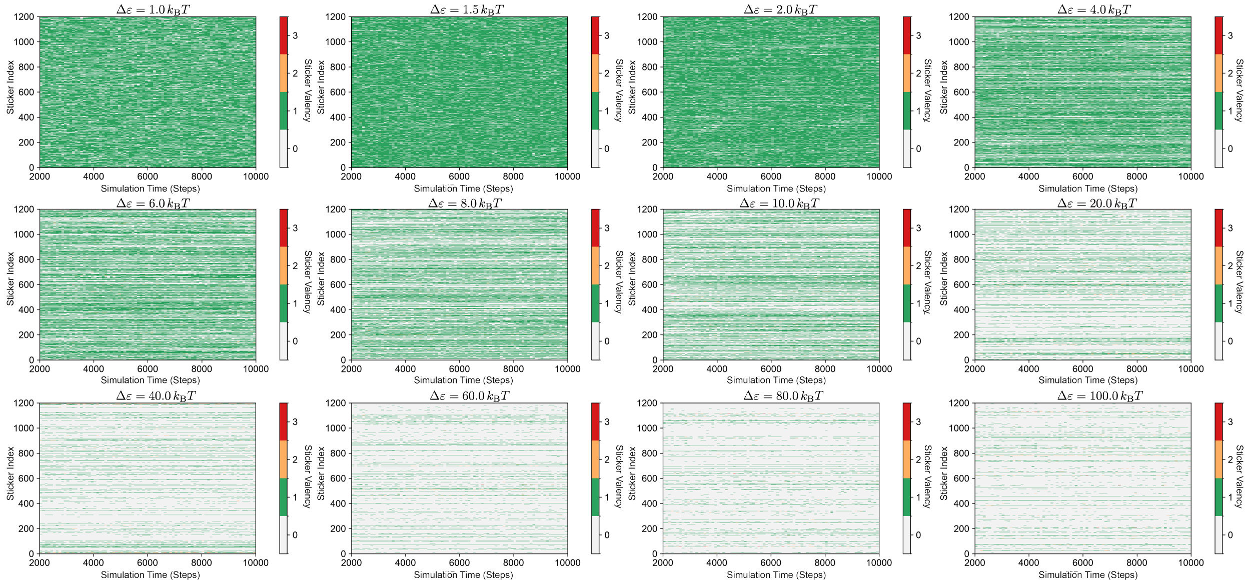


Figure S2: **Time-resolved sticker valency at different levels of spacer-interaction heterogeneity.** Each row corresponds to an individual sticker, indexed on the $y$-axis, and the $x$-axis denotes simulation time. Color indicates the instantaneous sticker valency, $v_i(t)$, defined as the number of other stickers bound to sticker $i$ at time $t$ as in Eq. 10 of the main text. The trajectories are the same as those analyzed in Figure 3B.

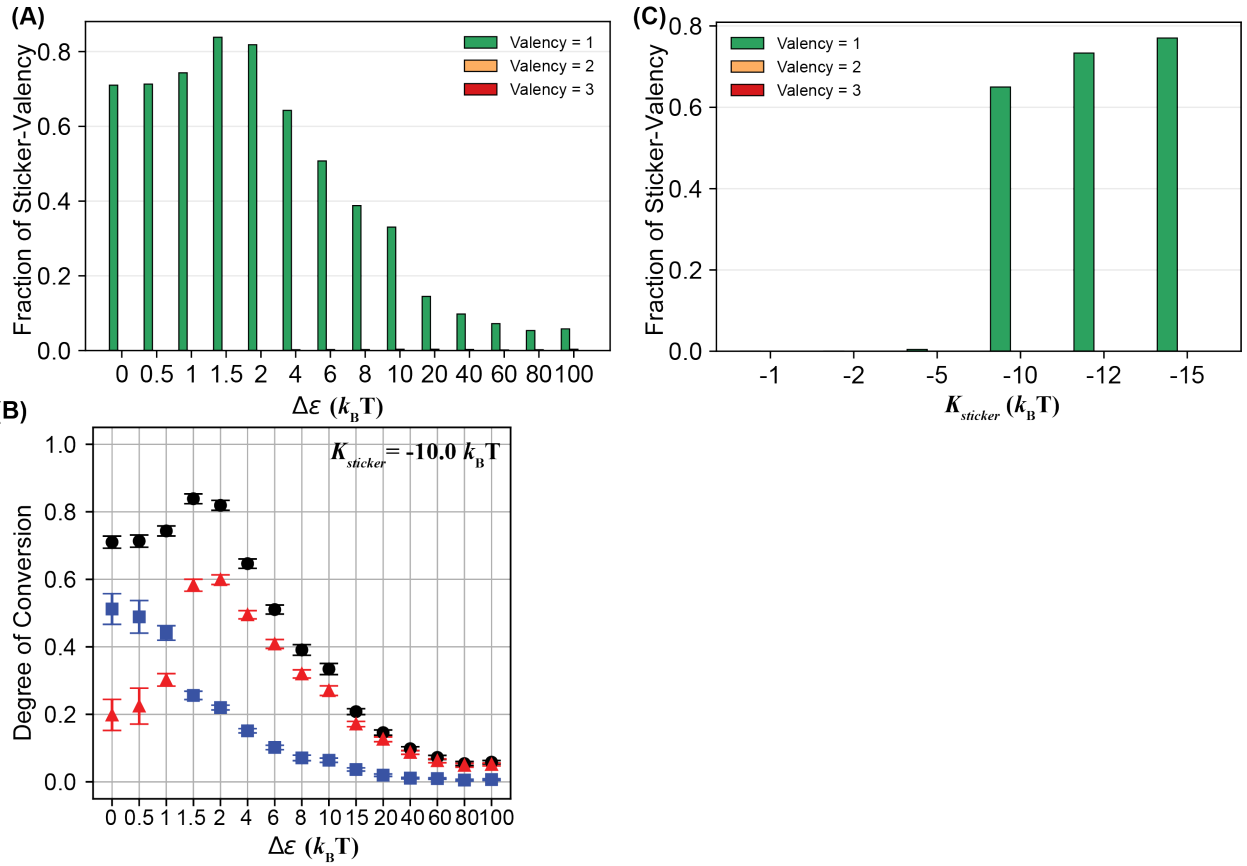


Figure S3: **Sticker valency statistics and degree of conversion for one-component systems.** (A) Fraction of bound sticker valency and (B) degree of conversion as a function of $\Delta\epsilon$, at fixed $K_{\text{sticker}} = -10\,k_{\text{B}}T$ and $\bar{\epsilon} = 0\,k_{\text{B}}T$. The fraction of unbound stickers (valency = 0) was not presented for clarity. (C) Fraction of sticker valency as a function of $K_{\text{sticker}}$ at fixed $\bar{\epsilon} = 0\,k_{\text{B}}T$ and $\Delta\epsilon = 0\,k_{\text{B}}T$. The corresponding degree of conversion analysis is shown in Figure 3A.

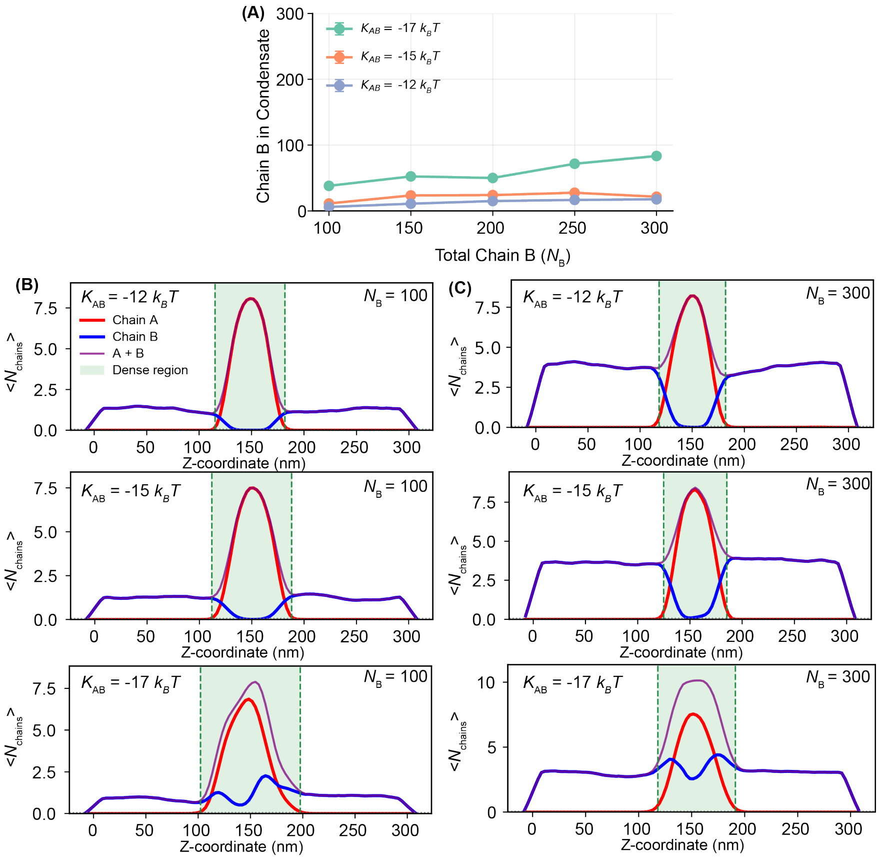


Figure S4: **The saturable recruitment observed in Figure 4A is robust to changes in sticker interaction strength.** (A) Number of B molecules recruited into the A condensate as a function of the total number of B molecules in the system for sticker-mediated A–B interactions. Simulations were performed using the same protocol as in Figure 4A but with different A–B sticker interaction strengths. The curve for $K_{\mathrm{AB}} = -17\, k_{\mathrm{B}}T$ is identical to that shown in Figure 4A. (B–C) Representative density profiles for systems with $N_{\mathrm{B}} = 100$ and $N_{\mathrm{B}} = 300$ at different A–B sticker interaction strengths.

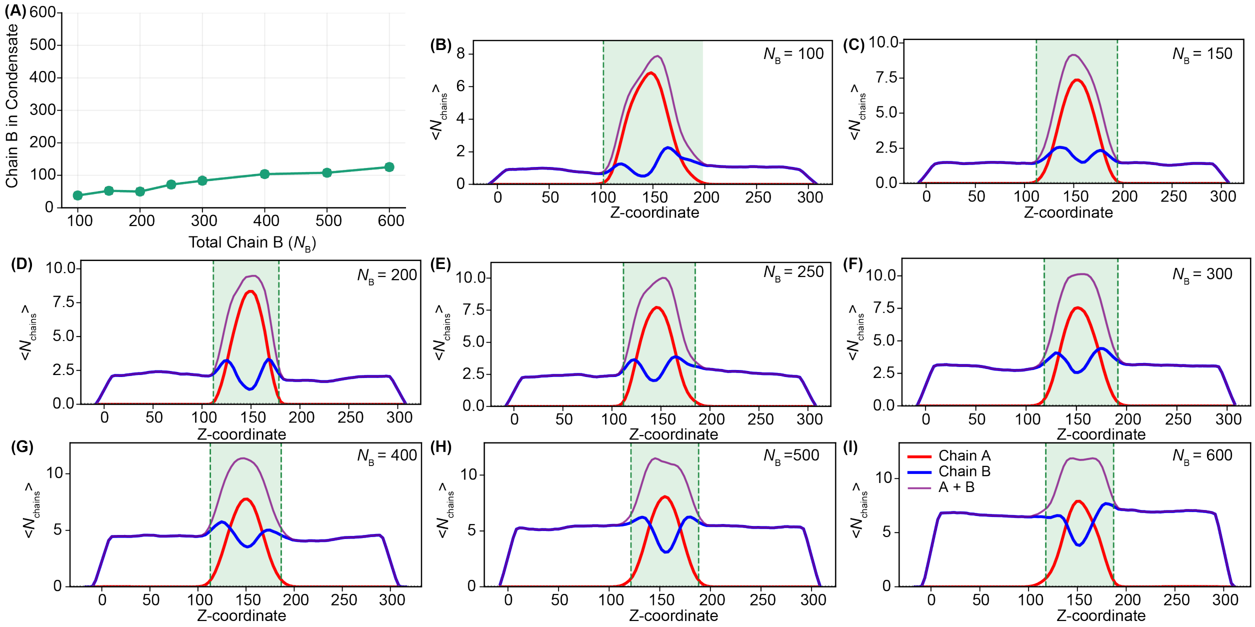


Figure S5: **Specific sticker interactions buffer client recruitment in two-component condensates.** (A) Number of B molecules recruited into the A condensate as a function of the total number of B molecules in the system for sticker-mediated A–B interactions. The data for $N_B \leq 300$ are presented in Figure 4A. (B-I) Density profiles for the simulations presented in panel A.

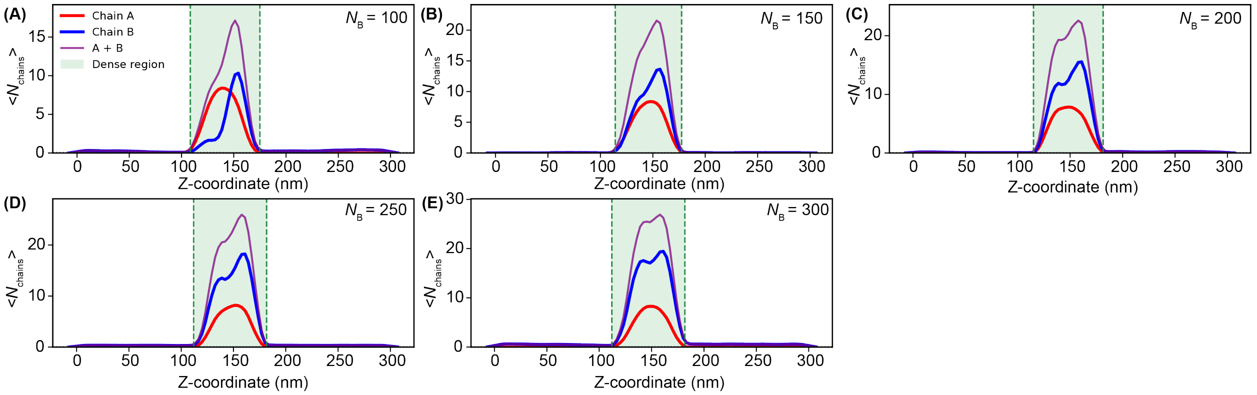


Figure S6: **(A-E) Density profiles for simulations presented in Figure 4C.**

Table S1: **Explicit architectures of chain A and chain B polymers.** All backbone beads are spacers. Auxiliary sticker beads are shown under their parent backbone bead. For A–A interactions and the one-component valency analysis, the default selector uses the even-numbered auxiliary stickers on Chain A, $S_2, S_4, \ldots, S_{24}$, corresponding to 12 active stickers per A chain. This default was used for the one- and two-component systems unless otherwise specified. For Figure 2B, the A–A selector was varied by `[::12]`,`[::6]`, `[::4]`, `[::3]`, `or [::2]` (every 12th, 6th, 4th, 3rd, or 2nd auxiliary sticker) to obtain $N_{\text{sticker}} = 2, 4, 6, 8,$ or 12 active stickers per Chain A. For A–B interactions, the complementary odd-numbered stickers on Chain A, $S_1, S_3, \ldots, S_{23}$, interact with all four sticker beads on Chain B.

**Chain A: spacer positions 1–25**

| Position | 1 | 2 | 3 | 4 | 5 | 6 | 7 | 8 | 9 | 10 | 11 | 12 | 13 | 14 | 15 | 16 | 17 | 18 | 19 | 20 | 21 | 22 | 23 | 24 | 25 |
|---|---|---|---|---|---|---|---|---|---|---|---|---|---|---|---|---|---|---|---|---|---|---|---|---|---|
| Spacer | $A_1$ | $A_2$ | $A_3$ | $A_4$ | $A_5$ | $A_6$ | $A_7$ | $A_8$ | $A_9$ | $A_{10}$ | $A_{11}$ | $A_{12}$ | $A_{13}$ | $A_{14}$ | $A_{15}$ | $A_{16}$ | $A_{17}$ | $A_{18}$ | $A_{19}$ | $A_{20}$ | $A_{21}$ | $A_{22}$ | $A_{23}$ | $A_{24}$ | $A_{25}$ |
| Auxiliary sticker | | | | $S_1$ | | | $S_2$ | | | $S_3$ | | | $S_4$ | | | $S_5$ | | | $S_6$ | | | $S_7$ | | | $S_8$ |
| Selected for A–A binding | | | | | | | $S_2$ | | | | | | $S_4$ | | | | | | $S_6$ | | | | | | $S_8$ |
| Selected for A–B binding (A side) | | | | $S_1$ | | | | | | $S_3$ | | | | | | $S_5$ | | | | | | $S_7$ | | | |

**Chain A: spacer positions 26–50**

| Position | 26 | 27 | 28 | 29 | 30 | 31 | 32 | 33 | 34 | 35 | 36 | 37 | 38 | 39 | 40 | 41 | 42 | 43 | 44 | 45 | 46 | 47 | 48 | 49 | 50 |
|---|---|---|---|---|---|---|---|---|---|---|---|---|---|---|---|---|---|---|---|---|---|---|---|---|---|
| Spacer | $A_{26}$ | $A_{27}$ | $A_{28}$ | $A_{29}$ | $A_{30}$ | $A_{31}$ | $A_{32}$ | $A_{33}$ | $A_{34}$ | $A_{35}$ | $A_{36}$ | $A_{37}$ | $A_{38}$ | $A_{39}$ | $A_{40}$ | $A_{41}$ | $A_{42}$ | $A_{43}$ | $A_{44}$ | $A_{45}$ | $A_{46}$ | $A_{47}$ | $A_{48}$ | $A_{49}$ | $A_{50}$ |
| Auxiliary sticker | | | $S_9$ | | | $S_{10}$ | | | $S_{11}$ | | | $S_{12}$ | | | $S_{13}$ | | | $S_{14}$ | | | $S_{15}$ | | | $S_{16}$ | |
| Selected for A–A binding | | | | | | $S_{10}$ | | | | | | $S_{12}$ | | | | | | $S_{14}$ | | | | | | $S_{16}$ | |
| Selected for A–B binding (A side) | | | $S_9$ | | | | | | $S_{11}$ | | | | | | $S_{13}$ | | | | | | $S_{15}$ | | | | |

**Chain A: spacer positions 51–74**

| Position | 51 | 52 | 53 | 54 | 55 | 56 | 57 | 58 | 59 | 60 | 61 | 62 | 63 | 64 | 65 | 66 | 67 | 68 | 69 | 70 | 71 | 72 | 73 | 74 |
|---|---|---|---|---|---|---|---|---|---|---|---|---|---|---|---|---|---|---|---|---|---|---|---|---|
| Spacer | $A_{51}$ | $A_{52}$ | $A_{53}$ | $A_{54}$ | $A_{55}$ | $A_{56}$ | $A_{57}$ | $A_{58}$ | $A_{59}$ | $A_{60}$ | $A_{61}$ | $A_{62}$ | $A_{63}$ | $A_{64}$ | $A_{65}$ | $A_{66}$ | $A_{67}$ | $A_{68}$ | $A_{69}$ | $A_{70}$ | $A_{71}$ | $A_{72}$ | $A_{73}$ | $A_{74}$ |
| Auxiliary sticker | | $S_{17}$ | | | $S_{18}$ | | | $S_{19}$ | | | $S_{20}$ | | | $S_{21}$ | | | $S_{22}$ | | | $S_{23}$ | | | $S_{24}$ | |
| Selected for A–A binding | | | | | $S_{18}$ | | | | | | $S_{20}$ | | | | | | $S_{22}$ | | | | | | $S_{24}$ | |
| Selected for A–B binding (A side) | | $S_{17}$ | | | | | | $S_{19}$ | | | | | | $S_{21}$ | | | | | | $S_{23}$ | | | | |

**Chain B: spacer positions 1–14**

| Position | 1 | 2 | 3 | 4 | 5 | 6 | 7 | 8 | 9 | 10 | 11 | 12 | 13 | 14 |
|---|---|---|---|---|---|---|---|---|---|---|---|---|---|---|
| Spacer | $B_1$ | $B_2$ | $B_3$ | $B_4$ | $B_5$ | $B_6$ | $B_7$ | $B_8$ | $B_9$ | $B_{10}$ | $B_{11}$ | $B_{12}$ | $B_{13}$ | $B_{14}$ |
| Auxiliary sticker | | | | $T_1$ | | | $T_2$ | | | $T_3$ | | | $T_4$ | |
| Selected for A–B binding (B side) | | | | $T_1$ | | | $T_2$ | | | $T_3$ | | | $T_4$ | |
